# A Systematic Review of the Biomechanical Effects of Harness and Head-Collar use in Dogs

**DOI:** 10.1101/759258

**Authors:** S Blake, R Williams, R Ferro de Godoy

## Abstract

The number of dogs in the UK is on the rise, as are canine sports involving the use of a harness to allow the dog to pull against an interface in the same way as a husky might pull a sled. Service dogs and those involved in essential work commonly wear a harness throughout their working lives, yet little is understood regarding the biomechanical impact of their use. This systematic review was conducted to review reported evidence of the biomechanical effects of harness and head collar (Halti) use in dogs.

Searches were applied covering 1910 to 2018 on the following databases: PubMed, Web of Science and Writtle Discovery.

Three publications were identified as suitable which were then critically evaluated using predefined criteria and ARRIVE based guidelines for bias assessment. Only one was considered to provide the most reliable data regarding the influence of harnesses on gait, whilst the remainder were considered to suffer a variety of issues including poor sample size, repeatability and study execution. The most appropriate study found that wearing a chest strap harness reduced shoulder extension in both walk and trot by up to 8° of movement, whilst a Y-shaped harness commonly marketed as non-restrictive reduced shoulder extension by up to 10° of movement, suggesting that the use of harness type restraints can affect canine gait, whereas no studies were found relating to the biomechanical effects of head-collar usage.

## Introduction

The canine population in the UK is currently estimated to be in excess of 9 million, whilst owner expenditure is in excess of £10 million per annum [1]. A fundamental requirement of dog ownership is control outside of the home, and owners spend even more time and money on puppy classes, obedience training and behaviourists in the hope of having a sociable and obedient pet, yet nearly a quarter of dogs given up to the Dogs Trust are there because of behavioural issues, such as a lack of control or aggression towards other dogs and/or humans [2].

A common solution for owners when faced with an unruly dog is the use of a restraint such as a harness or head collar (commonly known as a Halti), with manufacturers routinely advertising them on the basis of how they can benefit the owner, using product names such as Non-Pull™ and Easy walk™. Training a dog is vital in their early years and the foundation of correct behaviour [3] and harnesses are often used during the training period or as a training aid. It is surmised therefore that an owner is more likely to use these types of restraint when an animal is younger and relatively unruly, which raises questions regards their suitability and possible impact on a developing musculoskeletal system and its associated growth plates.

Canine sports such as Canicross (also known as Cani-fit) and Bikejoring are also growing in popularity in the UK, and these sports use harness systems to allow an animal to pull against an interface in much the same way as a husky may pull a sled, utilising the canines instinct to pull against pressure [3]. Harness systems of varying designs are also worn by all manner of service dogs, from guide dogs to search dogs and those involved with armed forces and policing.

It is clearly appropriate that a dog is under control at all times, for its own safety and the safety or others, yet there is very little discussion around the welfare consequences of using restraint devices, or whether they may prevent walking at the most natural, biomechanically efficient gait. As such they may have the potential to impact the dogs long term health and potentially compromise welfare.

If this proves to be true then the resultant costs may far out way any initial training expenditure needed to negate the need for restraint devices - the cost of veterinary care continues to rise, with insurers paying out on average £2 million per day for pet claims, an increase of nearly 56% in the last eight years [4]. The most common pet insurance claim is joint related, costing an average of over £450 [5] with the typical veterinary fee for a cruciate ligament repair being around £1,200, whilst a hip replacement costs in excess of £3,500 [6].

The most prevalent musculoskeletal disease in dogs are degenerative joint disease (DJD) and arthritis, with dysplasia, cruciate and patellar issues making up over 20% of the total number [7]. A further assumption could therefore be made that if harnesses do impact a dog’s natural gait, they may be a contributing factor in any of these conditions or could hasten the onset of any pathology that a dog may already suffer from.

It is relatively well known that if a dog’s gait is dysfunctional or impaired compensatory mechanisms will ensue [8] In the longer term this can lead to hypertrophy/atrophy of various muscle groups, as well as a myriad of musculoskeletal pathologies. Research by King [9] found that incorrect biomechanics will lead to loss of joint confirmation and function, in turn leading to abnormal wear, which can cause inflammation and arthritic conditions [8,10] DJD and arthritis are the two most common musculoskeletal issues seen in dogs, and whilst conditions such as elbow and hip dysplasia have strong conformational links, they may be exacerbated by additional restrictions in gait. [3,11,12]. Tendinopathy of the supraspinatus, infraspinatus, biceps and infraspinatus myopathy are some of the most frequent conditions diagnosed in performance dogs [3] all caused by varying degrees of micro and macro trauma and repetitive strain. Forelimb gait-related issues and lameness in active dogs is commonly as a result of medial shoulder syndrome (MSS) caused by repetitive micro trauma to multiple elements of the shoulder joint [13,14] leading to partial tears, dystrophic mineralization, chronic tenosynovitis, peritendinous adhesions and contractures [14] of the affected muscle. Cruciate ligament disease has its genesis within conformation, as well as strong causal links to obesity and immune mediated diseases [15] so as such may not be seen as a condition directly created by compensatory gait mechanisms, however as previously noted if forelimb stride is compromised in some way, this will lead to a change in the biomechanics of the whole animal [12] once again potentially creating adverse pressures in the caudal anatomy which may exacerbate or hasten any conditions that the dog may be predisposed to. The aim of this study therefore was to conduct a systematic review into the effects of common restraint systems on canine gait, by identifying existing research relating to restraint use and their effects, as well as analysis of the research quality. A further objective was to identify any links stated within the research to canine musculoskeletal pathologies.

## Materials and Methods

A systematic review protocol/research proposal was completed and submitted to Writtle University College in September 2018, along with a request for ethical approval and a full risk assessment. The research proposal was approved by Writtle University College ethics committee in October 2018 with approval number 98363809/2018.

The search terms set out in table 1 were used to identify all relevant research relating to animal studies. No control was specified in this instance as no description was deemed appropriate Table 1. PICO terms used in search criteria.

**Table 1.** PICO terms used in search criteria.

|  |  |
| --- | --- |
| Population | (dog* OR bitch* OR canine OR K9 OR husky* OR puppy* OR “canis lupus familiaris”<br>OR canid NOT dogmatic) |
| Intervention | (harness OR restraint* OR “head collar” OR “head-collar” OR halti OR “no pull” OR<br>“no-pull” OR “non-pull” OR “gentle leader” OR “julius-k9” OR dogmatic OR ruffwear)<br>OR “vest harness”) |
| Control | No control was specified |
| Outcomes | (kinematic* OR range of motion OR rom OR goniometry OR ground reaction force OR<br>grf OR pressure OR limb OR lameness OR gait OR stride length OR stride frequency OR<br>kinetics OR motion OR locomotion OR force OR “force plate” OR “video analysis”) |

Initial searches were applied in December 2018 to the PubMed database via the NCBI website (1910 – Dec 2018), the Web of Science database via the web of knowledge website (1969-Dec 2018) and the Writtle Discovery database via the eds.b.ebscohost website (1979 – Dec 2018). Potentially suitable papers were stored using Zotero reference management software to allow subsequent screening and removal of duplicates. After initial screening to remove duplicates the exclusion and inclusion criteria contained in table 2 was applied to both the title and abstracts.

**Table 2.**
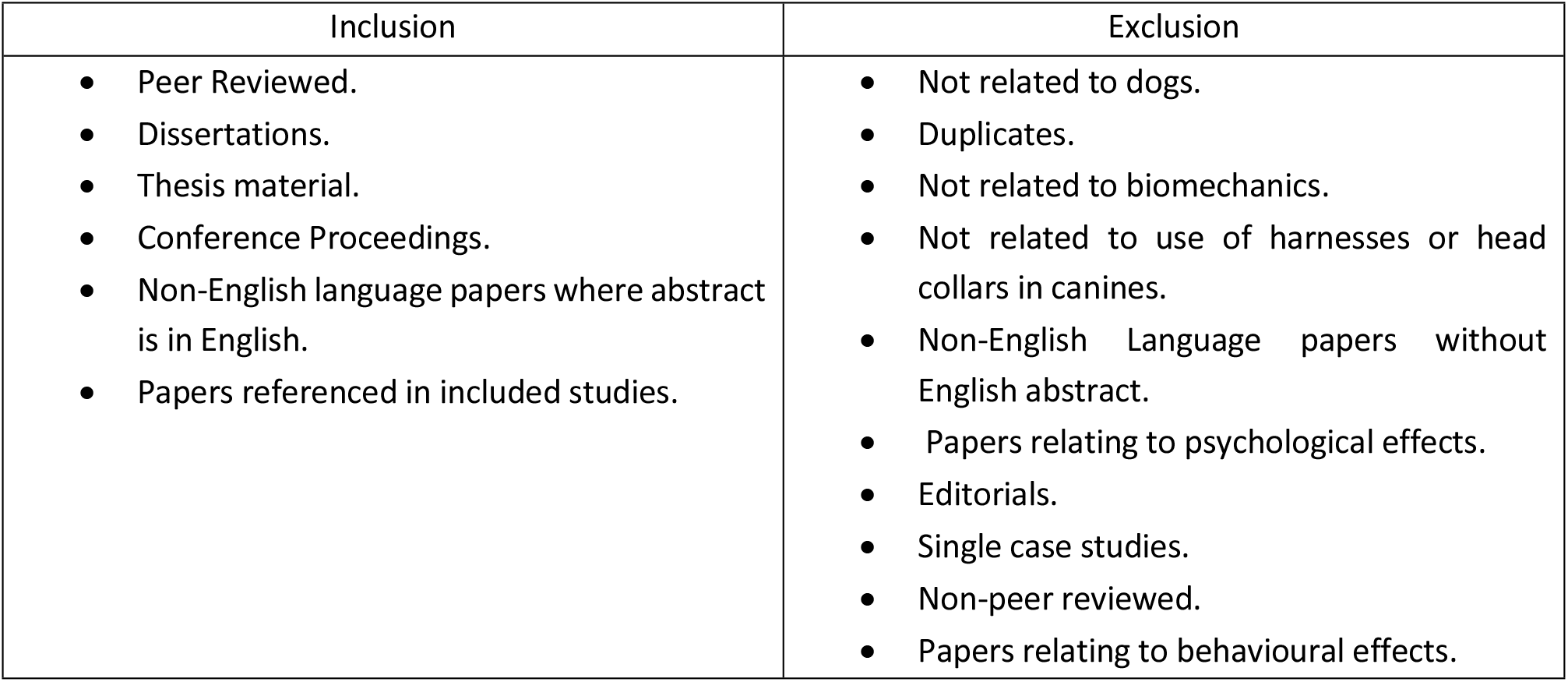
Inclusion and exclusion criteria.

The full text of any remaining papers was then used to confirm suitability. Bibliographies of the remaining papers were also used to identify any studies that were not located within the electronic search A standardised model of data collection was then used as set out within PRISMA guidelines [16] to extract key information from each of the included studies. Table 3 lists the relevant data that was included within the review.

**Table 3.** Data extracted from all papers deemed suitable for review

|  |
| --- |
| 1. Reference including publication date and author |
| 2. Study population |
| 3. Sample size |
| 4. Intervention |
| 5. Design |
| 6. Outcomes studied |
| 7. Main findings of the study |
| 8. Limitations of the study |

A bias assessment was conducted using ARRIVE (Animal Research: Reporting of *in vivo* experiments) guidelines 2018 to determine risk of bias. The full text of each paper was assessed as to whether it met the guidelines which would indicate a low risk of bias, did not meet the guidelines indicating a high risk of bias, or whether it partially met the guidelines indicating a medium risk of bias. Fourteen separate elements were considered for each study including study design, setting, study design reporting, procedures description, animal details, housing and husbandry, sample size, treatment allocation, outcome definition, statistical methods, baseline data, numbers analysed, outcomes and estimation and adverse events. All domains were then scored as either 1) low risk of bias 2) unclear risk of bias or 3) high risk of bias and results were collated using excel to produce a graph which would indicate the total risk of bias for the pool of papers as a whole.

In addition, papers included in the review were checked for evidence of conflicts of interest such as funding from organisations that may gain from specific research results.

## Results

Results of the search and subsequent exclusions can be seen in (Fig 1) whilst the results extracted from each study can be seen in table 4.

**Fig 1.**
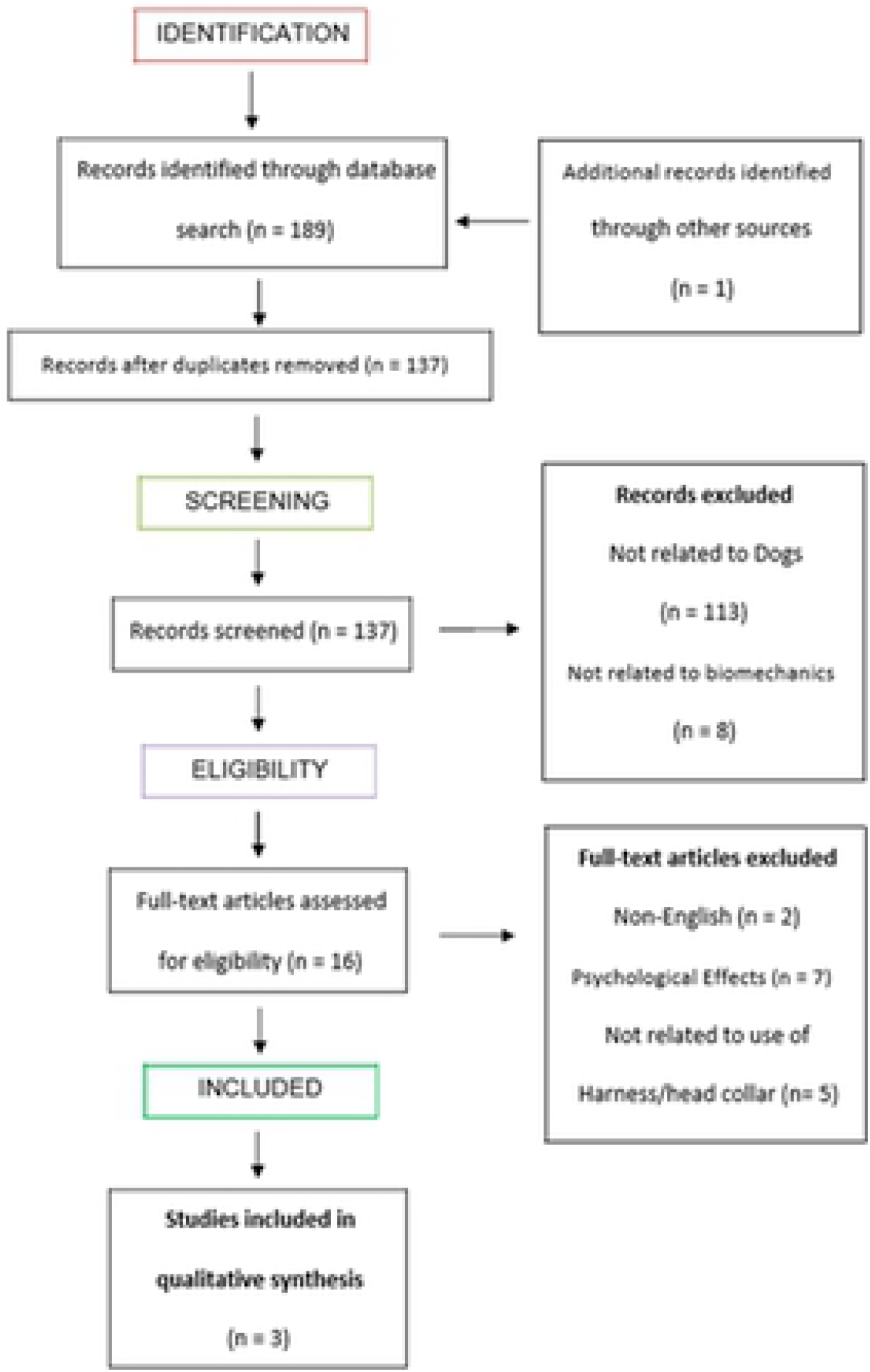
Flow chart to show search strategy used to identify articles regarding effects of harness and halter use on canine gait.

**Table 4.**
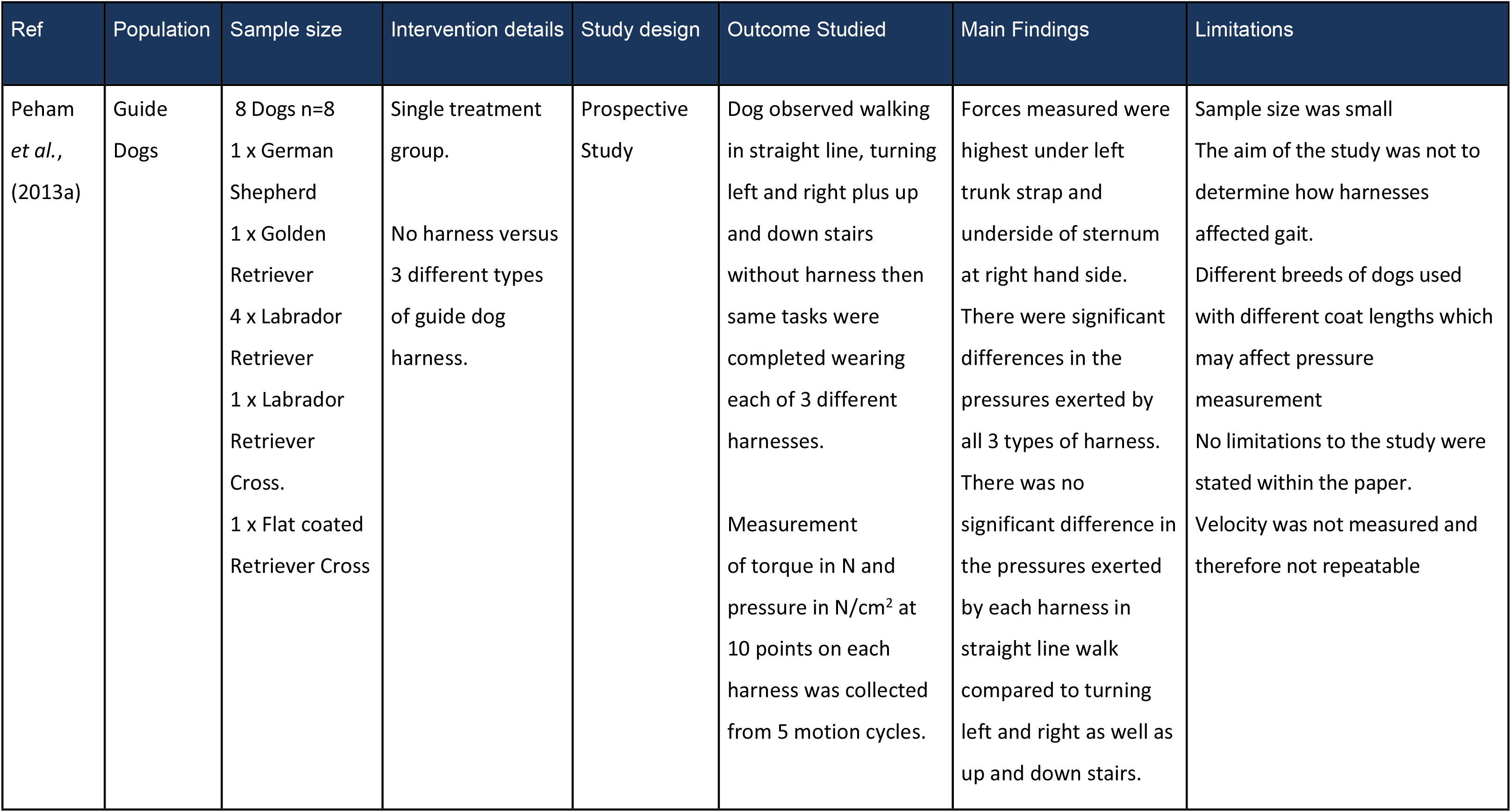
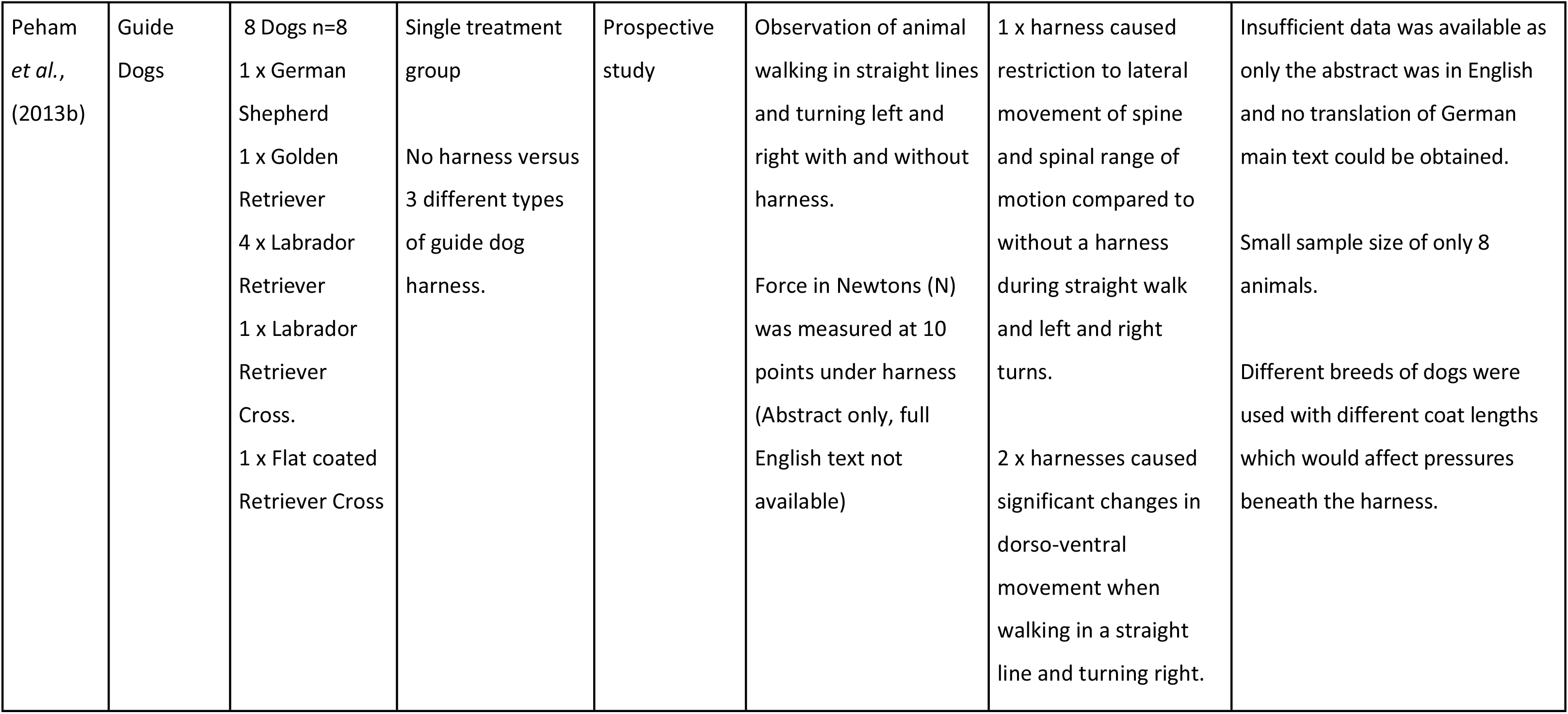
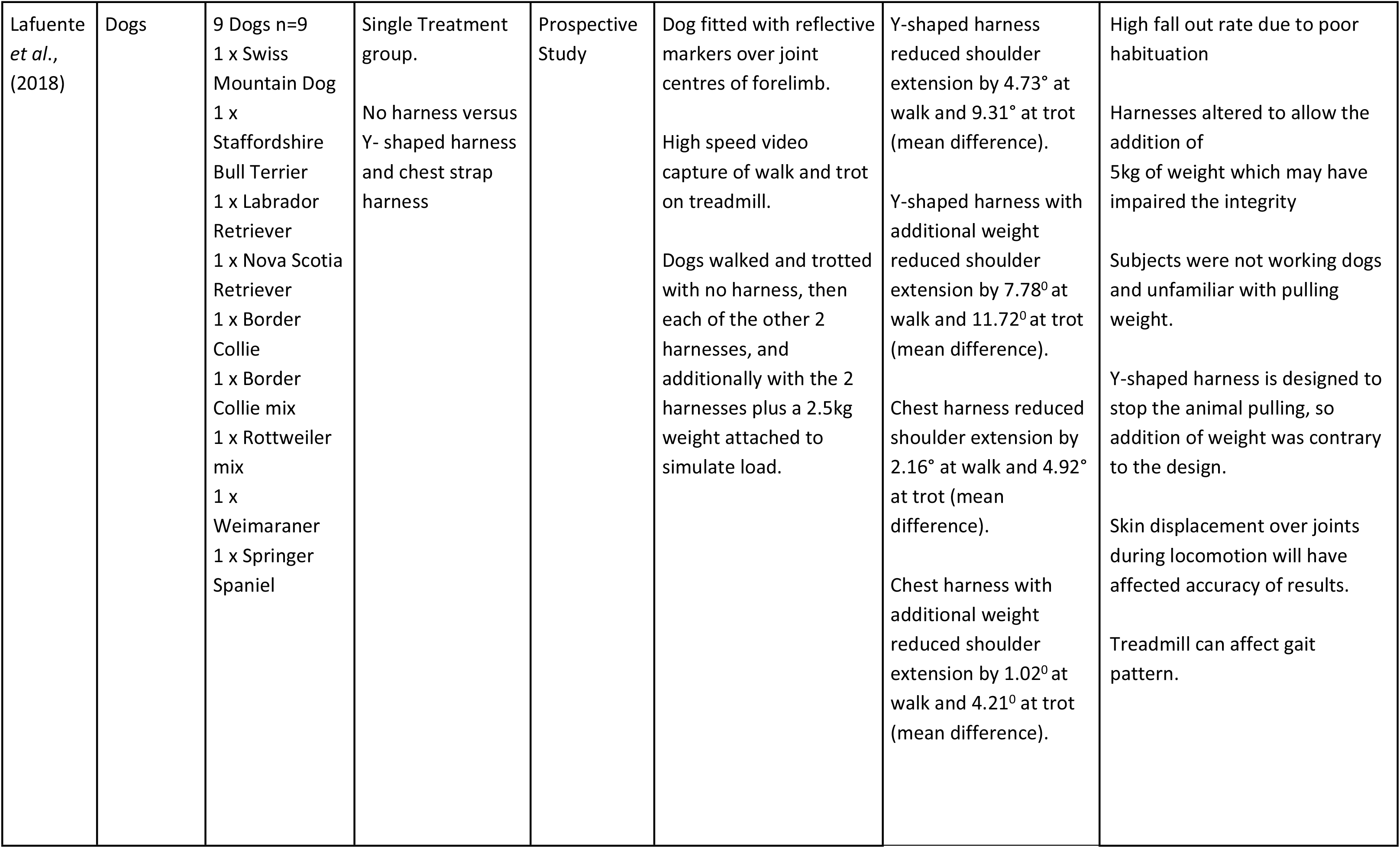
Results of Individual Studies

The three papers identified as suitable for review are as follows;

- Peham C, Limbeck S, Galla K, Bockstahler B. (2013a) Pressure distribution under three different types of harnesses used for guide dogs. The Veterinary Journal. (2013a);198: e93-e98 [17]
- Peham C, Limbeck S, Galla K, and Bockstahler B. Kinematic analysis of the influence of three different guide dog harnesses on the movement of the spine. Wiener Tierarztliche Monatsschrift.(2013b);100(11):306-312 [18]
- Lafuente M, Provis L, Schmalz E. Effects of restrictive and non-restrictive harnesses on shoulder extension in dogs at walk and trot. Veterinary Record. (2018):16-24 [19]

Lafuente *et al.* (2018) found that both a Y-shaped (non-restrictive) and chest harness restricted shoulder extension at both walk and trot, however the non-restrictive (Y-shaped) harness actually decreased shoulder extension more than the chest harness, by an additional 2.56° reduction in extension at walk and an additional 4.82° in trot. Full results are shown in table 5 and illustrated in (Fig 2).

**Fig 2.**
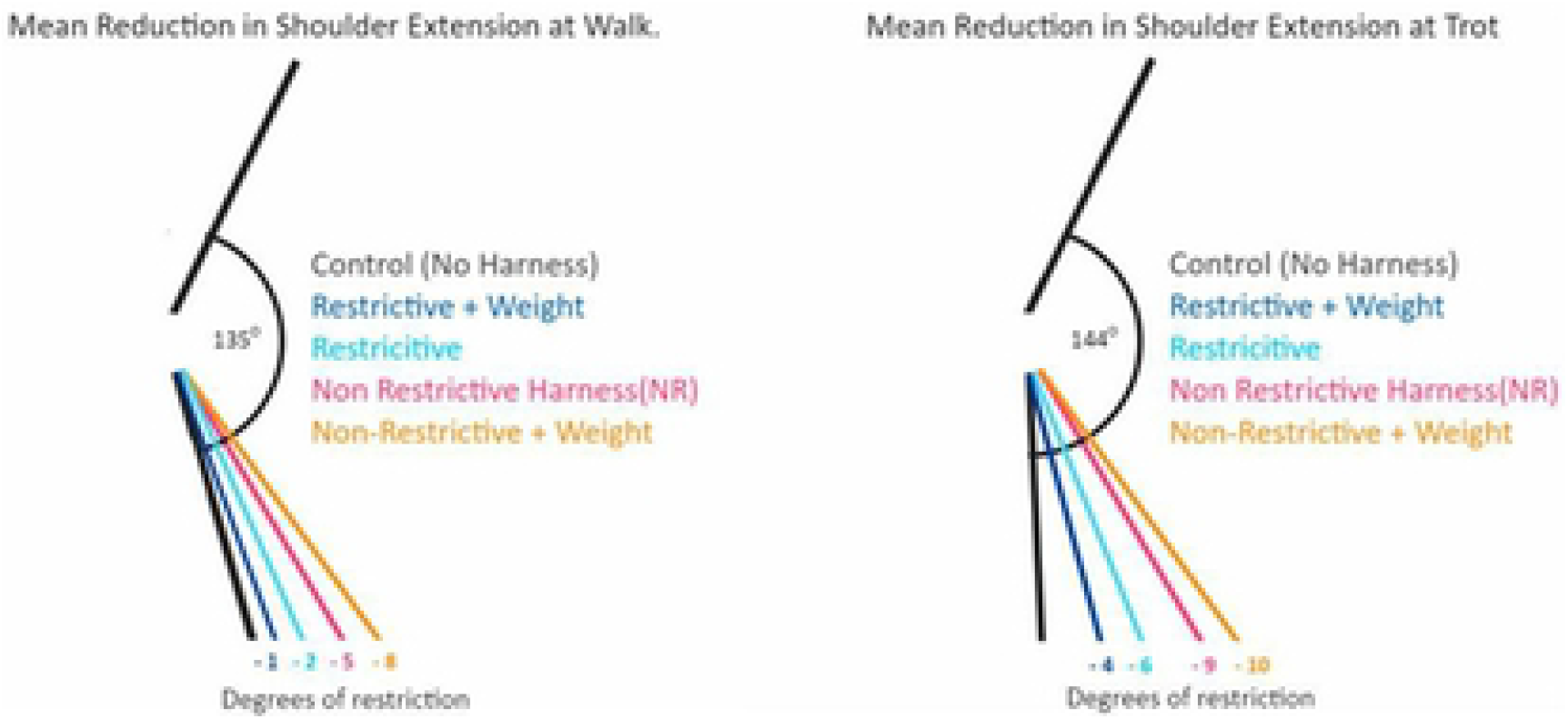
Reduction in mean shoulder extension in walk and trot, control versus non-restrictive harness (Y-shaped) and restrictive harness (chest harness). Adapted from Lafuente *et al.* (2018).

**Table 5.** Reduction in mean shoulder extension in walk and trot in degrees of movement, control versus Y Shaped and chest harness. Adapted from Lafuente *et al.* (2018).

| Harness | Walk (Degrees of movement) | Trot (Degrees of movement) |
| --- | --- | --- |
| Control | 135 ± 9.90 | 144 ± 8.38 |
| Y – Shaped Non-Restrictive | 130 ± 9.04 | 134 ± 11.69 |
| Non-Restrictive + 5 kg weight | 127 ± 11.00 | 133 ± 13.49 |
| Chest Harness | 133 ± 6.58 | 139 ± 9.71 |
| Chest Harness + 5 kg weight | 134 ± 6.82 | 140 ± 10.30 |

Peham *et al.* (2013a) found that force and pressures underneath all of the guide dog harnesses were highest at the right sternum, with both the left and right sternum constantly loaded by all three harnesses. There was insignificant loading of the spine from all three types of harnesses studied, as well as variable loading of the shoulders as seen in (Fig 4). Data from the study can be seen in table 6.

**Fig 3.**
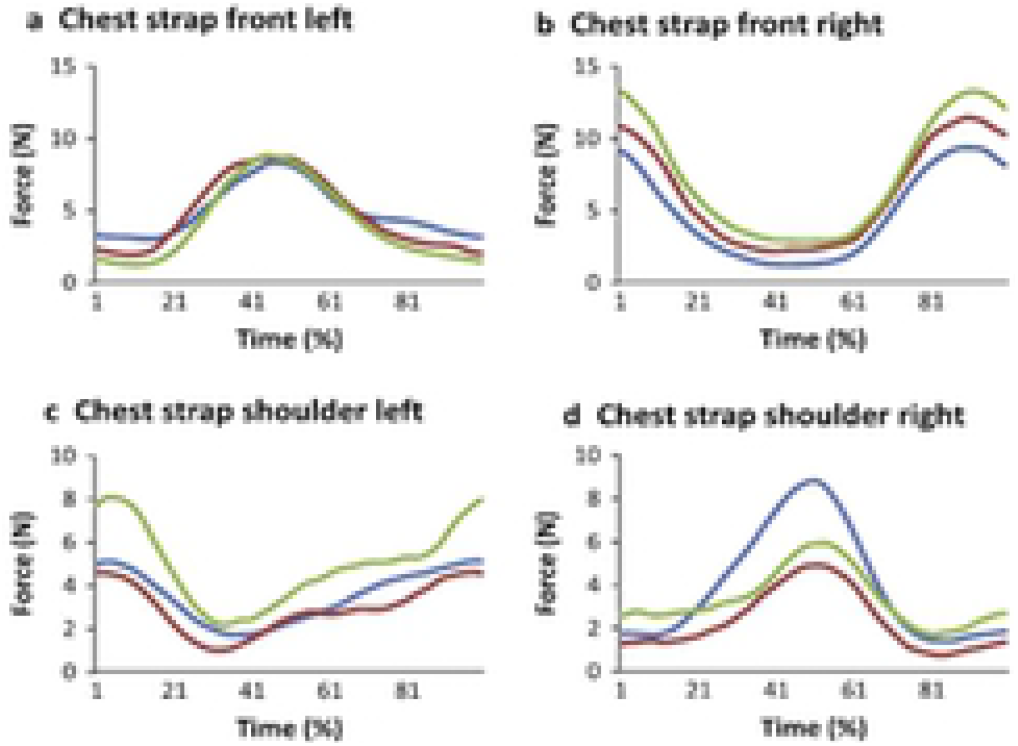
Force curves of chest strap regions of three guide dog harnesses during a straight walk exercise. (a) Chest strap left. (b) Chest strap right. (c) Chest strap shoulder left. (d) Chest strap shoulder right. Different harnesses represented by colour. Peham *et al.* (2013a)

**Figure 4.**
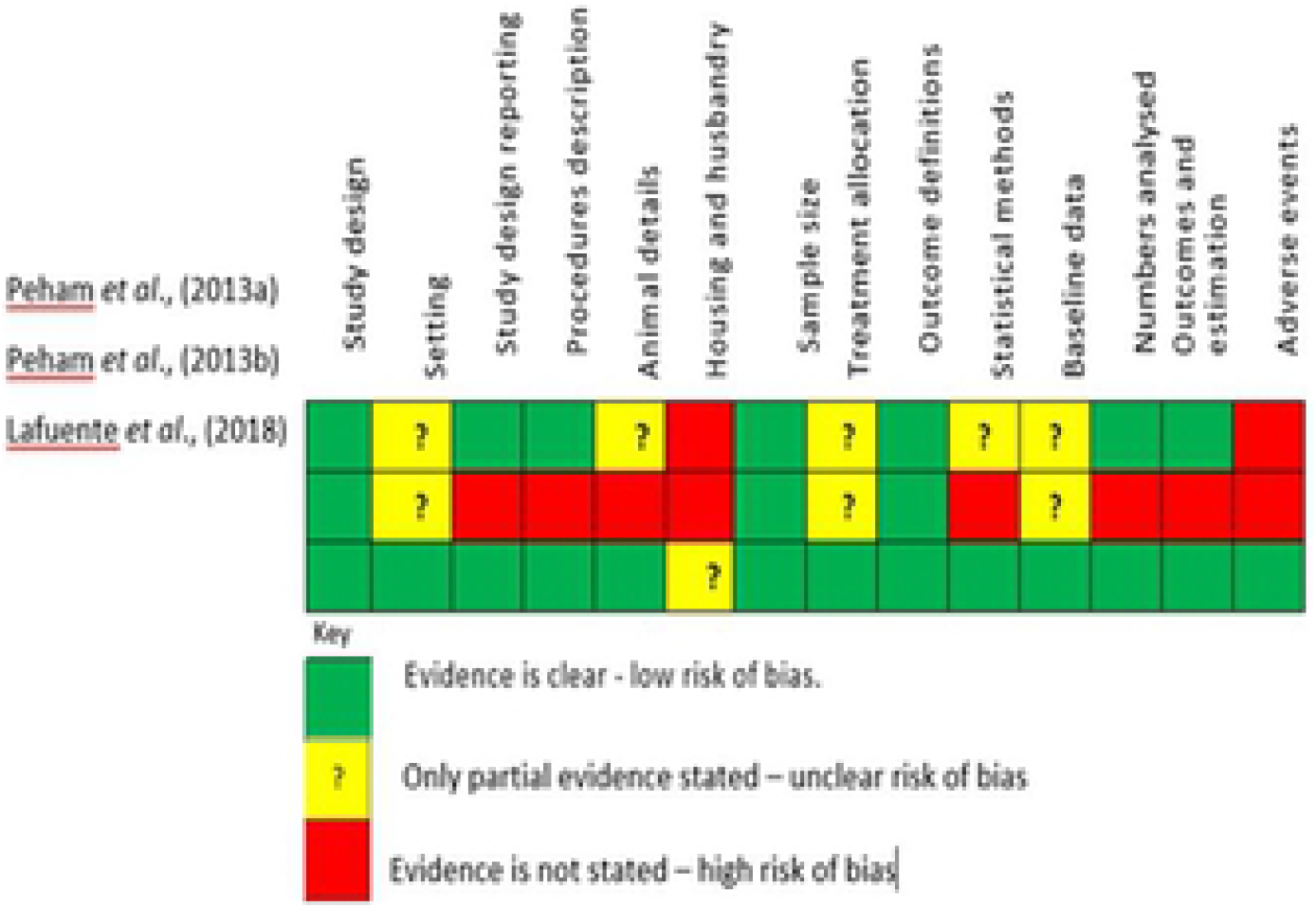
Results of bias analysis

**Table 6:**
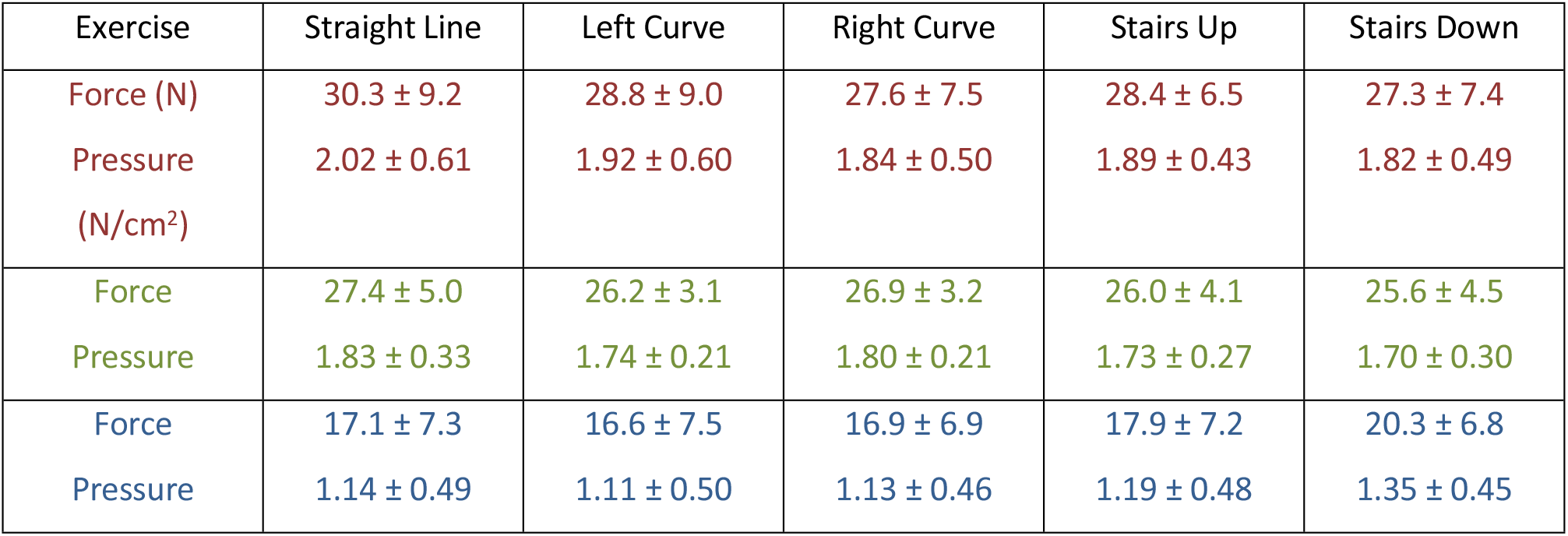
Different chest strap harnesses represented by colours corresponding to figure 16. Adapted from Peham *et al.* (2013a).

| Exercise | Straight Line | Left Curve | Right Curve | Stairs Up | Stairs Down |
| --- | --- | --- | --- | --- | --- |
| Force (N) | 30.3 ± 9.2 | 28.8 ± 9.0 | 27.6 ± 7.5 | 28.4 ± 6.5 | 27.3 ± 7.4 |
| Pressure (N/cm<sup>2</sup>) | 2.02 ± 0.61 | 1.92 ± 0.60 | 1.84 ± 0.50 | 1.89 ± 0.43 | 1.82 ± 0.49 |
| Force | 27.4 ± 5.0 | 26.2 ± 3.1 | 26.9 ± 3.2 | 26.0 ± 4.1 | 25.6 ± 4.5 |
| Pressure | 1.83 ± 0.33 | 1.74 ± 0.21 | 1.80 ± 0.21 | 1.73 ± 0.27 | 1.70 ± 0.30 |
| Force | 17.1 ± 7.3 | 16.6 ± 7.5 | 16.9 ± 6.9 | 17.9 ± 7.2 | 20.3 ± 6.8 |
| Pressure | 1.14 ± 0.49 | 1.11 ± 0.50 | 1.13 ± 0.46 | 1.19 ± 0.48 | 1.35 ± 0.45 |

The second study by Peham *et al.*,(2013b) only reported data via an abstract which states that one harness restricted “latero-lateral motion of the spine, causing a significant restricted minimum and maximum lateral movement and ROM” whilst the same harness plus one other caused “significant changes in the dorso-ventral movement of the spine”. A summary of publications and their results can be seen in table 7.

**Table 7.**
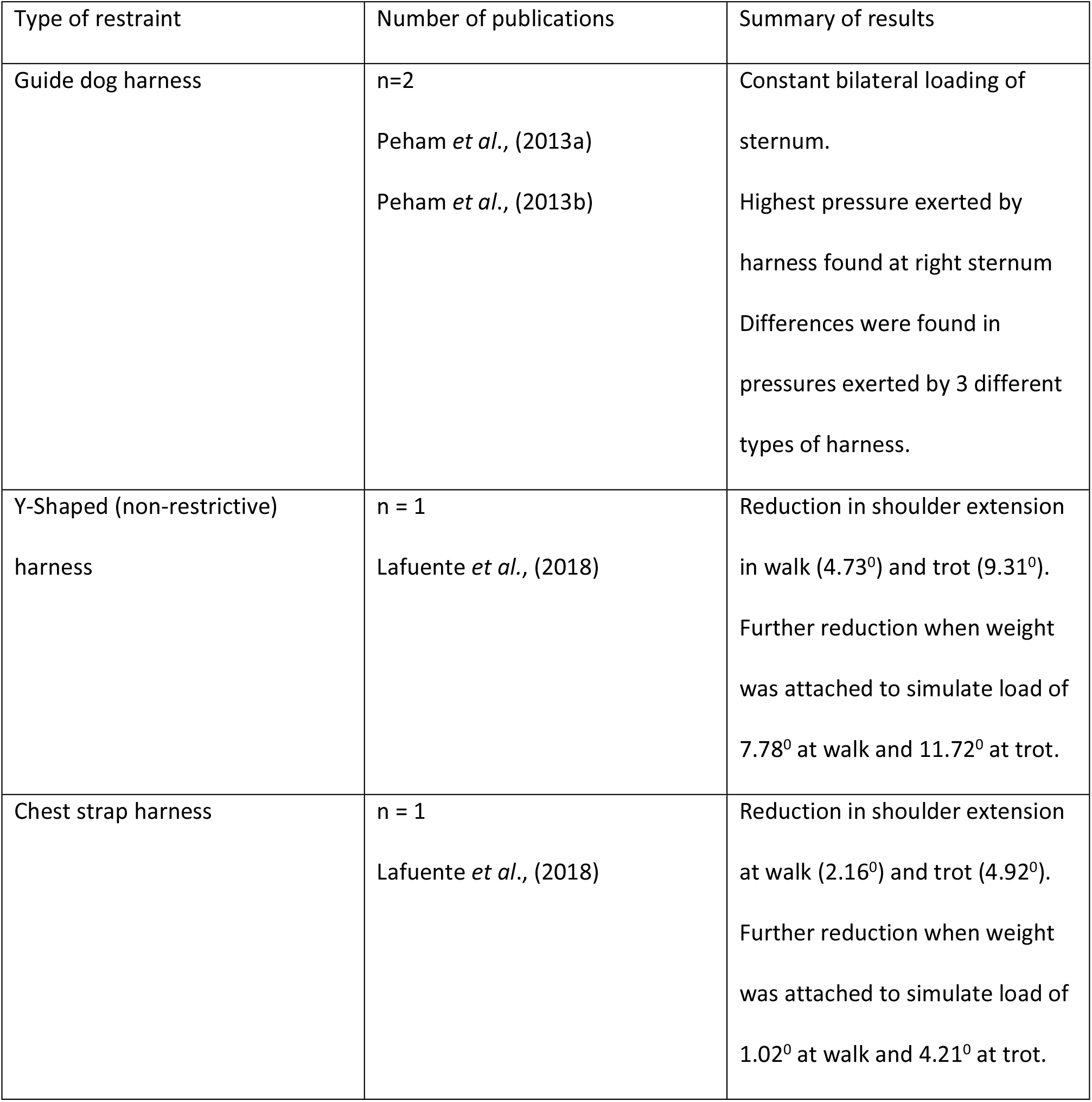
Summary of publications and results.

| Type of restraint | Number of publications | Summary of results |
| --- | --- | --- |
| Guide dog harness | n=2<br>Peham *et al.*, (2013a)<br>Peham *et al.*, (2013b) | Constant bilateral loading of sternum.<br>Highest pressure exerted by harness found at right sternum<br>Differences were found in pressures exerted by 3 different types of harness. |
| Y-Shaped (non-restrictive) harness | n = 1<br>Lafuente *et al.*, (2018) | Reduction in shoulder extension in walk (4.73°) and trot (9.31°).<br>Further reduction when weight was attached to simulate load of 7.78° at walk and 11.72° at trot. |
| Chest strap harness | n = 1<br>Lafuente *et al.*, (2018) | Reduction in shoulder extension at walk (2.16°) and trot (4.92°).<br>Further reduction when weight was attached to simulate load of 1.02° at walk and 4.21° at trot. |

### Bias Assessment

The individual results of the ARRIVE Bias assessment of included studies are shown in (Fig 4). No conflicts of interest were Identified. Two papers failed to report details of animal housing and husbandry including procedures to monitor test subject’s welfare during the study, whilst the remaining publication partially disclosed husbandry only. Two papers failed to fully discuss how treatments were allocated to each test subject, although as they were cohort studies no randomisation was expected. Two papers also did not fully disclose baseline data and as previously mentioned only the Lafuente *et al.,* (2018) study measured velocity which would allow a comparison with baseline measurements. Risk of bias is necessary when discussing validity of results and overall it is felt that the above limitations do not affect the validity of the data.

## Discussion

Although not conclusive it is clear that harnesses utilising a chest strap or of a Y-shaped design do limit the angle of shoulder extension at both walk and trot. The reasons why a Y-shaped harness, deemed non-restrictive would limit extension to a greater degree is unclear, however the author postulates that it may restrict the musculature around the scapula at both the cranial angle and border which would reduce its extension. It is also unclear whether the width of strap or padding would further influence angulation, although a reduction in the width of straps would focus pressure beneath them. Only the Lafuente *et al.* (2018) study specified a width of 25 millimetres for the Y-shaped harness straps, running from the sternum to the dorsal neck so no conclusions can be drawn regarding width of straps versus the effect on gait, but is worthy of further study as it could be of detriment to the dog if areas are constantly loaded or at areas of high pressure such as the sternum as indicated with the Peham *et al.* 2013a study. The Lafuente *et al.* (2018) study used two 2.5kg weights attached to the lead on either side of the dog to simulate pulling and interestingly this addition reduced shoulder extension even further. This was not consistent with both types of harness, indicating that the shape of the harness could be a contributing factor as opposed to the load pulling the limb caudally. It may also be that the dog shifts its centre of mass cranially to allow it to pull more effectively, which is especially pertinent where canine sports such as canicross are concerned as the animal is expected to be able to manoeuvre at speed, with a harness that is padded enough so as not to cause injury, but thin enough to allow the limbs to move freely. The addition of 5 kg of weight is relatively light when compared to the potential forces caused by a runner attached via a bungee lead, especially if the lead is at the end of its stretch capacity. One unexpected result is that a guide dog harness did not create pressure on the dorsal spine, but this may be due to the handler needing to maintain contact by lifting the harness slightly via the handle. This would also explain why forces are highest at the right sternum as guide dogs are taught to walk on the right of their owner at all times, meaning a slight lifting force would be exerted by the handler from the left side of the dog. It is assumed that the majority of dog owners do not use a harness handle when exercising their pet so the slight shift in weight bearing would not be significant for the wider canine population, however it does have implications if a dog is undergoing therapy that requires them to use a sling device or harness, in that an additional load will be created on the limb opposite the handler, and therefore the handler should be on the same side as any affected limb so as not to place additional strain on the area. Albeit this particular study did not interpret its results in terms of gait, it did show that the forces involved are relatively high, even at walk, at just over 30N. Studies on the effects of poor tack fitting in equines have found that a similar force can cause dry spots under the saddle, indicative of skin atrophy as the sweat glands within the capillaries have been damaged [20]. A dog’s mass is smaller than that of a horse, meaning the exerted load is higher in relation to their mass than that experienced by equines. Further research would be needed to ascertain whether the same could be true of harnesses used in canine sports, but what is known is that ischemic damage can occur quickly when skin is put under pressure [21] and by the time damage is noticeable the underlying muscles will also have been affected, as skin tissue is generally the last tissue to show signs of macroscopic damage [20]. A dog’s coat will also make it much more difficult to spot evidence of ischemic damage but conversely may act as a form of padding to reduce overall pressures and potential tissue damage. Although only limited information was available from the Peham *et al.* (2013b) study into spinal movement it did conclude that a harness will impact lateral movement of the spine which adds an additional dimension – a harness will need to allow for adequate flexion and extension, lateral bending, and axial rotation of the spine, all of which will alter through changes in head and neck position at different gaits. It would therefore seem logical that the larger or wider the harness, the more these will be impaired. As has been noted skin displacement over anatomical landmarks during locomotion can lead to incorrect data collection [22] so this would also need to be addressed in futures studies. No studies to date have explored any impact on gait when using a head collar and leash, which would be necessary if it is to be compared to the suitability of harnesses.

Risk of bias was low, but none of the studies adequately discussed the housing and husbandry of the test subjects, and almost all did not fully examine or record baseline data prior to any intervention which again limits the validity of the results.

### Strength of Evidence and Further Research

All of the studies are limited by the small number of animals taking part. Harris *et al.* [23] suggested a sample size of at least 27 subjects is needed to be able to collect clinically relevant data, which could be problematic for further research. The Lafuente *et al.* (2018) study did start with a sample of 30 dogs but could only collect data on nine due to most being unfamiliar with the treadmill. Some research does exist with regards the amount of time a dog may need to become habituated to its study environment, and the data within the above studies suggested that dogs became comfortable with use of a treadmill after 30 minutes, whilst a study on greyhounds suggested useful data may be gathered in as little as 30 seconds [24] possibly due to a greyhounds natural inclination to run, or familiarity with training expectations. Another study by Rumph et al. [25] found that poor habituation impacts hind limb stance times and lower impulses of vertical force. Speed of forward motion will be influenced by stride length and limb angulation, yet only the Lafuente *et al.* (2018) study included a reliable measure of velocity, which is vital if further researchers wish to build on what is already known. Any research is also going to be hampered by the huge variance in breeds, conformations and even gaits – there is no such thing as a “typical” canine so a strategy could be to study one breed in particular first where it could be most useful, for example a working dog breed such as Spaniels commonly used for search work. Interestingly the chest type harness is commonly used for these types of dog, as such it needs to allow full ROM of the forelimb, and if gait is affected performance and working longevity may suffer. Recommendations for further research are therefore myriad – it would seem logical to assess the impact of restraint systems on dogs most at risk of harm through daily use such as those used by policing and security services as mentioned above. This would also reduce the overall number of breeds that would need to be studied initially as well as having the greatest potential impact. What is clear is that future studies will need to be of a sufficiently robust nature to be able to provide appropriate data, which has been lacking in some of the research so far.

The only clinically relevant data that can be taken from this review is that shoulder extension is limited by two of the most common types of harness. At present the use of relatively low-cost technology to assess gait is still underutilised in veterinary practice, but what is clear is that quantitative analysis is the most effective way of detecting biomechanical abnormalities as well as the underlying reasons.

## Conclusion

As has been shown very little research exists regarding the effect of restraint use on canine gait. Of the studies identified only one would be deemed to have the necessary scientific protocols to show sufficient evidence of a change in gait, however it lacks a large enough sample size to reflect on the canine population as a whole. None of the studies showed a biomechanical change when using a head halter but questions do remain as to their long-term suitability. Nor does current research relate to any forms of pathology which would be the next logical step, otherwise as standalone research the value is limited. This lack of understanding poses a dilemma for veterinarians and physiotherapists alike, especially in the context of evidence-based practice, who are forced to make judgements on what is best for a dog’s long-term welfare, with no reliable means of knowing potential outcomes.

Further research is needed to establish if limiting a dog’s natural gait impacts their longer-term welfare and to define the relationship between certain types of harness and injury, especially in working breeds and those taking part in sporting endeavours. Owners, veterinarians and physiotherapists need to understand the importance of the correct selection of a canine restraint system based on the breed as well as the dog’s purpose. Special consideration should be given to working dogs and they may routinely have to adopt an abnormal gait, as well as canine athletes who may be subject to the same restrictions but also be expected to work at their maximum capacities.

## References

1. Pet Food Manufacturing Association. [Internet] London: c2018 [cited 2018 1 Dec]. Available at: https://www.pfma.org.uk/ [cited 2018 1 Dec].

2. Dogs Trust [Internet] London:c2018 [cited 2018 18 Nov]. Available at: https://www.dogstrust.org.uk/about-us/

3. Ogburn P, Crouse S, Martin F, Houpt K. Comparison of behavioural and physiological responses of dogs wearing two different types of collars. Applied Animal Behaviour Science. 1998 61(2):133–142.

4. ABI [Internet]. London: Pet insurance claims data; c2017 [cited 2018 November 26]. Available from: https://www.abi.org.uk/news/news-articles/2018/05/pet-claims-are-through-the-woof

5. This is Money. [Internet]. London; c2018 [cited 2018 November 26]. Available from: https://www.thisismoney.co.uk/money/bills/article-3533744/Vet-fees-hit-810-average-uninsured-pets-treatments-cost-8k.html.

6. Animal Trust [Internet]. Ellesmere Port: [cited 2018 November 26]. Vet Treatment Prices; c2017 [cited 2018 November 26]. Available from: https://www.animaltrust.org.uk/prices

7. RVC. [Internet]. London; c2018 [cited 2018 November 26]. Available from: https://www.rvc.ac.uk/vetcompass/learn-zone/infographics/canine

8. Impellizer J, Tetrick M, Muir P. Effect of weight reduction on clinical signs of lameness in dogs with hip osteoarthritis. Journal of the American Veterinary Medical Association.2000; 216(7):1089–1091

9. King, M. Etiopathogenesis of Canine Hip Dysplasia, Prevalence, and Genetics. Veterinary Clinics of North America: Small Animal Practice. 2017; 47(4):53–767

10. Guilak F, Ratcliffe A, Lane N, Rosenwasser M, Mow V. Mechanical and biochemical changes in the superficial zone of articular cartilage in canine experimental osteoarthritis. Journal of Orthopaedic Research. 1994;12(4):474–484

11. Preston T, Wills A. A single hydrotherapy session increases range of motion and stride length in Labrador retrievers diagnosed with elbow dysplasia. The Veterinary Journal. 2018; 234:105–110

12. Smith G, Mayhew P, Kapatkin A, McKelvie P, Shofer F, Gregor T. Evaluation of risk factors for degenerative joint disease associated with hip dysplasia in German Shepherd Dogs, Golden Retrievers, Labrador Retrievers, and Rottweilers. Journal of the American Veterinary Medical Association. 2001;219(12):1719–1724

13. Kunkel K, Rochat M. A Review of Lameness Attributable to the Shoulder in the Dog: Part Two. Journal of the American Animal Hospital Association. 2008;44(4):163–170

14. Marcellin-Little D, Levine D, Canapp S. The Canine Shoulder: Selected Disorders and Their Management with Physical Therapy. Clinical Techniques in Small Animal Practice. 2007;22(4):171–182

15. Zink C, Carr B. Locomotion and Athletic Performance. Canine Sports Medicine and Rehabilitation.2018: 23–42

16. Moher D, Liberati A, Tetzlaff J, Altman D. Preferred Reporting Items for Systematic Reviews and MetaAnalyses: The PRISMA Statement. Journal of Clinical Epidemiology. 2009;62(10):1006–1012

17. Peham C, Limbeck S, Galla K, Bockstahler B. Pressure distribution under three different types of harnesses used for guide dogs. The Veterinary Journal. 2013;198: e93–e98

18. Peham C, Limbeck S, Galla K, Bockstahler B. Kinematic analysis of the influence of three different guide dog harnesses on the movement of the spine. Wiener Tierarztliche Monatsschrift. 2013; 100(11):306–312

19. Lafuente M, Provis L, Schmalz E. Effects of restrictive and non-restrictive harnesses on shoulder extension in dogs at walk and trot. Veterinary Record.2018;18–24

20. Von Peinen K, Wiestner T, Von Rechenberg B, Weishaupt M. Relationship between saddle pressure measurements and clinical signs of saddle soreness at the withers. Equine Veterinary Journal. 2010;42: 650–653

21. Clayton H, O’Connor K, Kaiser L. Force and pressure distribution beneath a conventional dressage saddle and a treeless dressage saddle with panels. The Veterinary Journal. 2014;199(1):44–48

22. Filipe V, Pereira J, Costa L, Maurício A, Couto P, Melo-Pinto P, et al. Effect of skin movement on the analysis of hindlimb kinematics during treadmill locomotion in rats. Journal of Neuroscience Methods.2006; 153(1):55–61

23. Harris K, Whay R, Murrell C. An investigation of mechanical nociceptive thresholds in dogs with hind limb joint pain compared to healthy control dogs. The Veterinary Journal, 2018;234(4):85–90.

24. Richards J, Clements D, Drew S, Bennett D, Carmichael S, Owen M. Kinematics of the elbow and stifle joints in greyhounds during treadmill trotting. Veterinary and Comparative Orthopaedics and Traumatology. 2004;17(03):141–145

25. Rumph P, Steiss J, Montgomery R. Effects of selection and habituation on vertical ground reaction force in Greyhounds. American Journal Veterinary Medicine. 1997; 58:1206–1208

